# Using open data to rapidly benchmark biomolecular simulations: Phospholipid conformational dynamics

**DOI:** 10.1101/2020.11.09.374850

**Authors:** Hanne S. Antila, Tiago M. Ferreira, O. H. Samuli Ollila, Markus S. Miettinen

**Affiliations:** Department of Theory and Bio-Systems, Max Planck Institute of Colloids and Interfaces, 14424 Potsdam, Germany; NMR Group — Institute for Physics, Martin-Luther University Halle–Wittenberg, 06120 Halle (Saale), Germany; Institute of Biotechnology, University of Helsinki, 00014 Helsinki, Finland

## Abstract

Molecular dynamics (MD) simulations are widely used to monitor time-resolved motions of biomacromolecules, although it often remains unknown how closely the conformational dynamics correspond to those occurring in real life. Here, we used a large set of open-access MD trajectories of phosphatidylcholine (PC) lipid bilayers to benchmark the conformational dynamics in several contemporary MD models (force fields) against nuclear magnetic resonance (NMR) data available in the literature: effective correlation times and spin-lattice relaxation rates.

We found none of the tested MD models to fully reproduce the conformational dynamics. That said, the dynamics in CHARMM36 and Slipids are more realistic than in the Amber Lipid14, OPLS-based MacRog, and GROMOS-based Berger force fields, whose sampling of the glycerol backbone conformations is too slow. The performance of CHARMM36 persists when cholesterol is added to the bilayer, and when the hydration level is reduced. However, for conformational dynamics of the PC headgroup, both with and without cholesterol, Slipids provides the most realistic description, because CHARMM36 overestimates the relative weight of ~1-ns processes in the headgroup dynamics.

We stress that not a single new simulation was run for the present work. This demonstrates the worth of open-access MD trajectory databanks for the indispensable step of any serious MD study: Benchmarking the available force fields. We believe this proof of principle will inspire other novel applications of MD trajectory databanks, and thus aid in developing biomolecular MD simulations into a true computational microscope—not only for lipid membranes, but for all biomacromolecular systems.

## 1 Introduction

Ever since the conception of Protein Data Bank (PDB)^1,2^ and GenBank,^3,4^ open access to standardised and searchable pools of experimental data has revolutionized scientific research. Constantly growing and improving in fidelity due to collaborative effort,^5–8^ the now hundreds of databanks ^9^ fuel the data-driven development of biomolecular structure determination,^10^ refinement, ^11^ prediction,^12^ and design^13^ approaches, as well as development of drugs, ^14,15^ materials,^16,17^ and more.^18,19^ It is clear that open data enables scientific progress that is far beyond the resources of a single research group or institute. Consequently, the call for public availability and conservation of data has extended to molecular dynamics (MD) simulation trajectories of biomolecules,^20–22^ and the discussion on how and by whom such databanks for dynamic structures would be set up is currently active.^23–26^ While there are currently no general MD databanks in operation, individual databanks are accepting contributions on nucleic acid, ^27^ protein/DNA/RNA, ^28^ cyclodextrin, ^29^ G-protein-coupled receptor, ^30^ and lipid bilayer^31^ simulations.

Since 2013, the NMRlipids Project (nmrlipids.blogspot.fi) has promoted a fully open collaboration approach, where the whole scientific research process—from initial ideas and discussions to analysis methods, data, and publications—is all the time publicly available. ^32^ While its main focus has been on conformational ensembles of different lipid headgroups and on ion binding to lipid membranes,^32–34^ the NMRlipids Project has also built a databank^31^ (zenodo.org/communities/nmrlipids) containing hundreds of atomistic MD trajectories of lipid bilayers and indexed at `nmrlipids.fi`.

MD databanks are expected to be particularly relevant for disordered biomolecules, such as biological lipids composing cellular membranes or intrinsically disordered proteins. These, in contrast to folded proteins or DNA strands, cannot be meaningfully described by the coordinates of a single structure alone. Realistic MD simulations, however, can provide the complete conformational ensemble and dynamics of such molecules, as well as enable studies of their biological functions in complex biomolecular assemblies. Unfortunately, the current MD force fields largely fail to capture the conformational ensembles of lipid headgroups and disordered proteins. ^32,34-37^ Therefore, before they can be used to draw conclusions, the quality of MD simulations must always be carefully assessed against structurally sensitive experiments. For lipid bilayers, such evaluation is possible against NMR and scattering data.^38^

Here, we demonstrate the use of a preexisting, publicly available set of MD trajectories to rapidly evaluate the fidelity of phospholipid conformational dynamics in state-of-the-art force fields. The rate at which individual molecules sample their conformational ensemble is traditionally used to assess if a given MD simulation has converged. Going beyond such practicalities, realistic dynamics are particularly desired for the intuitive interpretation of NMR experiments sensitive to molecular motions, ^39^ as well as to understand the dynamics of biological processes where molecular deformations play a rate-limiting role, such as membrane fusion. ^40^ The here presented comprehensive comparison of dynamics between experiments and different MD models at various biologically relevant compositions and conditions is thus likely to facilitate the development of increasingly realistic phospholipid force fields.

Above all, our results demonstrate the power of publicly available MD trajectories in creating new knowledge at a lowered computational cost and high potential for automation. We believe that this paves the way for novel applications of MD trajectory databanks, as well as underlines their usefulness—not only for lipid membranes, but for all biomolecular systems.

## 2 Methods

### Lipid conformational dynamics in NMR data

We analyzed the veracity of phosphatidylcholine (PC) lipid dynamics in MD based on two quantities that are readily available from published^39,41-43 13^C-NMR experiments and directly quantifiable from atomistic MD simulations: The effective C–H bond correlation times *τ*_e_, and the spin-lattice relaxation rates *R*_1_.

### Effective C–H bond correlation times *τ*_e_

In a lipid bilayer in liquid crystalline state, each individual lipid samples its internal conformational ensemble and rotates around the membrane normal. Lipid conformational dynamics are reflected in the second order autocorrelation functions of its C–H bonds

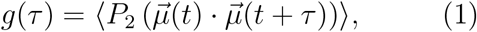

where the angular brackets depict time average, 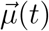 is the unit vector in the direction of the C–H bond at time *t*, and *P*_2_ is the second order Legendre polynomial 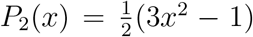. To analyze the internal dynamics of lipids, the C–H bond autocorrelation function is often written as a product

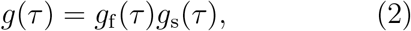

where *g*_f_(*τ*) characterizes the fast decays owing to, e.g., the internal dynamics and rotation around membrane normal, and *g*_s_(*τ*) the slow decays that originate from, e.g., lipid diffusion between lamellae with different orientations and periodic motions due to magic angle spinning conditions (Fig. 1). Ferreira et al. ^41^ have experimentally demonstrated that for all phospholipid carbons the motional correlation times contributing to *g*_f_ are well below *μ*s, and to *g*_s_ well above 100 *μ*s. This separation of time scales gives rise to the plateau 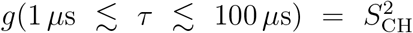 illustrated in Fig. 1. *S*_CH_ is the C-H bond order parameter

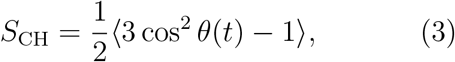

where *θ*(*t*) is the angle between the C–H bond and the bilayer normal. *S*_CH_ can be independently measured using dipolar coupling in ^13^C or quadrupolar coupling in ^2^H-NMR experiments. Knowing the set of *S_CH_* for all the C–H bonds in a lipid is highly useful in order to eval uate its conformational ensemble.^38^

**Figure 1:**
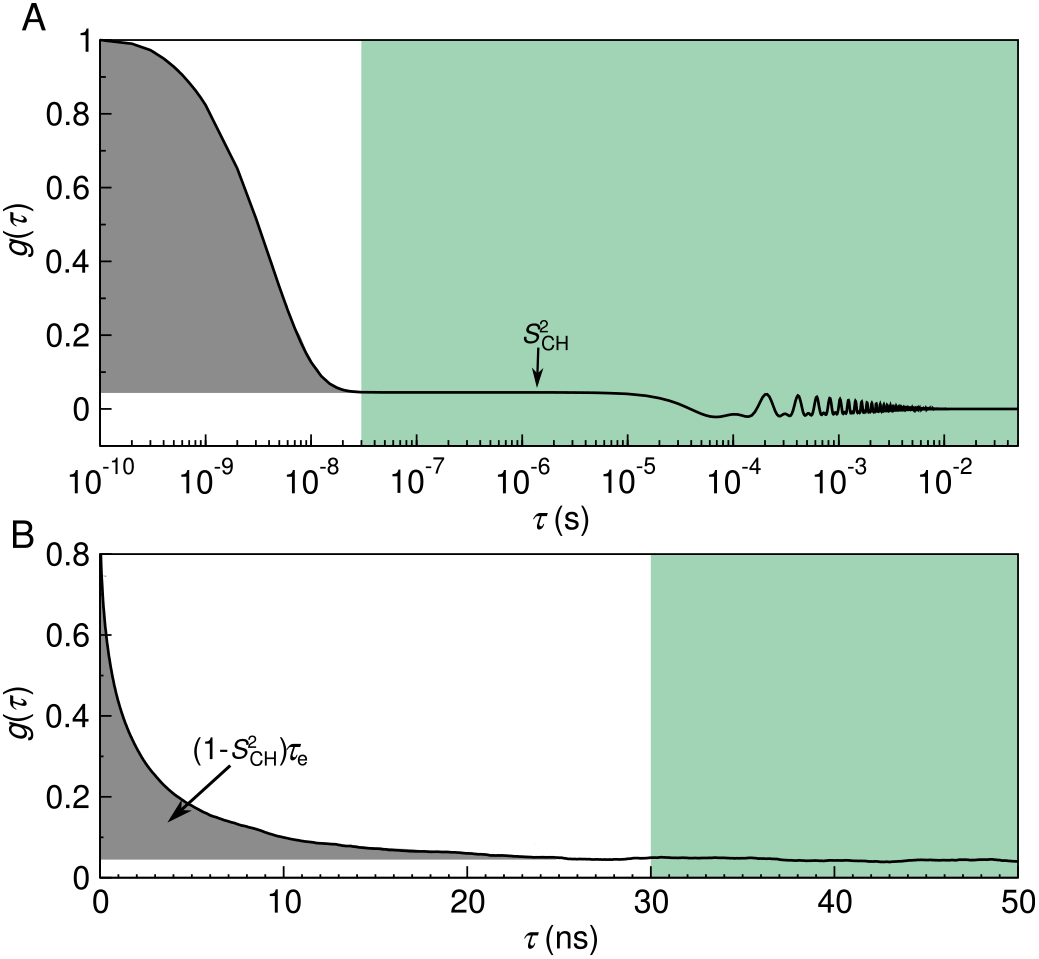
C–H bond autocorrelation function *g*(*τ*). (A) Idealised illustration of the fast (white background) and the slow (green) mode of the correlation function in solid-state NMR experiments. The fast mode decays to a plateau on which 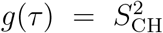, while the slow mode gives the final descent to zero. Oscillations at the slow mode region are due to magic angle spinning. (B) Typical *g*(*τ*) obtained from an MD simulation, showing the decay towards 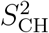. The gray area under the curve is equal to 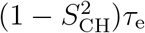.

As *S*_CH_ describe the conformational ensemble of the lipid, the fast-decaying component *g*_f_ of the C–H bond autocorrelation function intuitively reflects the time needed to sample these conformations. The complex internal dynamics containing multiple timescales can be conveniently summarized using the effective correlation time

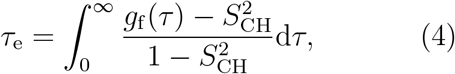

which is related to the gray-shaded area below the correlation function in Fig. 1. The *τ*_e_ detect essentially an average over all the time scales relevant for the lipid conformational dynamics. Their relation to process speeds is intuitive: Increase of long-lived correlations increases *τ*_e_.

### Spin-lattice relaxation rates *R*_1_

The C–H bond dynamics relate to *R*_1_, the spinlattice relaxation rate, through

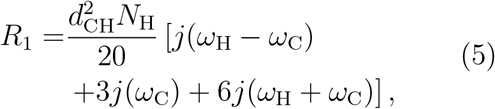

where *ω*_H_ is the ^1^H and *ω*_C_ the ^13^C-NMR Larmor frequency, and *N*_H_ the number of hydrogens covalently bonded to the carbon. The rigid dipolar coupling constant *d*_CH_ ≈ — 2π × 22kHz for the methylene bond. The spectral density *j*(*ω*) is given by the Fourier transformation

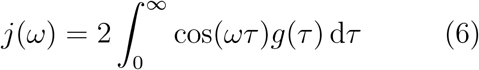

of the C–H bond autocorrelation function *g*(*τ*) (Eq. (1)). Clearly the connection between *R*_1_ and molecular dynamics is not straightforward; the magnitude of *R*_1_ does, however, reflect the relative significance of processes with timescales near the inverse of *ω*_H_ and *ω*_C_. These two frequencies depend on the field strength used in the NMR experiments: Typically *R*_1_ is most sensitive to motions with time scales ~0.1–10ns. (In our experimental data^39,41-43^ *ω*_C_ = 125 MHz and *ω*_H_ = 500 MHz, which gives (2π × 125 MHz)^-1^ = 1.3 ns and (2π × 625 MHz)^-1^ = 0.25ns.) A change in given *R*_1_, therefore, indicates a change in the relative amount of processes occurring in a window around the sensitive timescale; inferring also the direction to which the processes changed (speedup/slowdown) requires measuring *R*_1_ at various field strengths.

### Data acquisition and analysis

All the experimental quantities used in this work were collected from the literature sources^39,41-43^ cited at the respective figures.

The simulation trajectories were collected from the general-purpose open-access repository Zenodo (zenodo.org), with the majority of the data originating from the NMR-lipids Project^32,33^ (nmrlipids.blogspot.fi). The trajectories were chosen by hand based on how well the simulation conditions matched the available experimental data (lipid type, temperature, cholesterol content, hydration), and how precisely one could extract the quantities of interest from the trajectory (length of simulation, system size). Note that apart from the sampling accuracy, simulation size does not affect *τ*_e_ and *R*_1_ (Fig. 2). Table 1 lists the chosen trajectories of pure POPC (1-palmitoyl-2-oleoyl-glycero-3-phosphocholine) bilayers at/near room temperature and at full hydration; Table 2 lists the trajectories with cholesterol; and Table 3 those with varying hydration. Full computational details for each simulation are available at the cited Zenodo entry.

**Figure 2:**
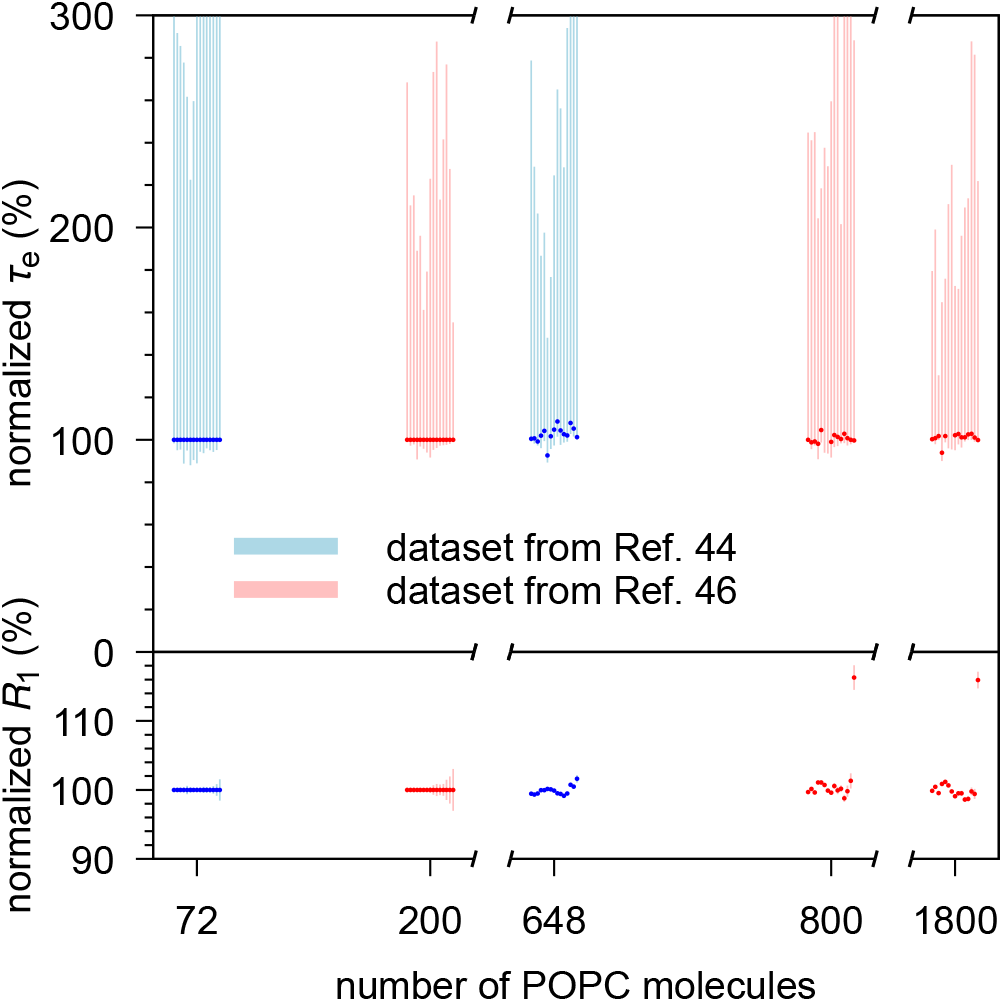
Effective correlation times (*τ*_e_, upper panel) and *R*_1_ rates (lower panel) do not markedly depend on the system size. Shown are two CHARMM36 POPC data sets that varied size while keeping other simulation parameters fixed: Ref. 44 (blue, system sizes 72 and 648 lipids)^45^ and Ref. 46 (red, system sizes 200, 800, and 1800 lipids). Both data sets are shown normalized against their smallest system. The 15 datapoints shown for each system correspond, from left to right, to the carbon segments *γ, β, α*, *g*_3_,…, C17/C15’, C18/C16’, cf. Fig. 3.

**Table 1:** Analyzed open-access MD trajectories of pure POPC lipid bilayers at full hydration. Note that the temperature varied across these openly available simulation data, but in no case was *T* lower than in the experiment. Thus, as dynamics slows down when temperature drops, any overestimation of *τ*_e_ by MD (as typically seen in Fig. 3) would get worse if the simulations were done at the experimental 298 K.

| force field lipid/water | $N_l^a$ | $N_w^b$ | $T^c$(K) | $t_{\text{anal}}^d$(ns) | files<sup>e</sup> |
| --- | --- | --- | --- | --- | --- |
| Berger-POPC-07<sup>47</sup>/SPC<sup>48</sup> | 256 | 10240 | 300 | 300 | [49] |
| CHARMM36<sup>50</sup>/TIP3P<sup>51</sup> | 256 | 8704 | 300 | 300 | [52] |
| MacRog<sup>53</sup>/TIP3P<sup>54</sup> | 128 | 5120 | 300 | 500 | [55] |
| Lipid14<sup>56</sup>/TIP3P<sup>54</sup> | 72 | 2234 | 303 | 50 | [57] |
| Slipids<sup>58</sup>/TIP3P<sup>54</sup> | 200 | 9000 | 310 | 500 | [59] |
| ECC<sup>60</sup>/SPC-E<sup>61</sup> | 128 | 6400 | 300 | 300 | [62] |

<sup>a</sup>Number of POPC molecules.
<sup>b</sup>Number of water molecules.
<sup>c</sup>Simulation temperature.
<sup>d</sup>Trajectory length used for analysis.
<sup>e</sup>Reference for the openly available simulation files.

**Table 2:**
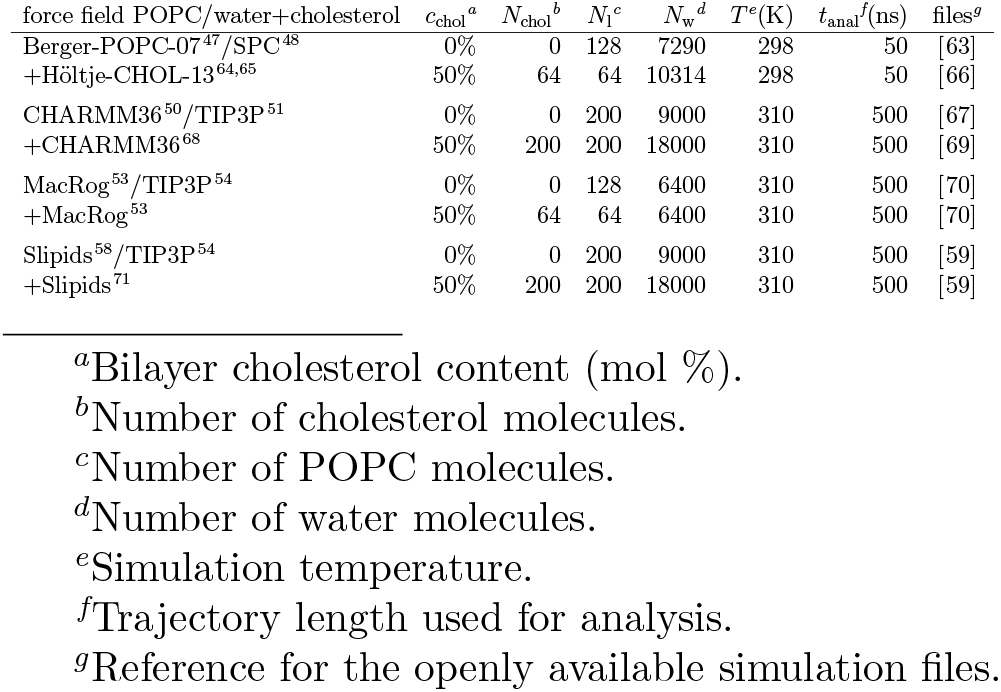
Analyzed open-access MD trajectories of cholesterol-containing POPC bilayers at full hydration.

| force field POPC/water+cholesterol | $c_{\text{chol}}^a$ | $N_{\text{chol}}^b$ | $N_l^c$ | $N_w^d$ | $T^e$(K) | $t_{\text{anal}}^f$(ns) | files<sup>g</sup> |
| --- | --- | --- | --- | --- | --- | --- | --- |
| Berger-POPC-07<sup>47</sup>/SPC<sup>48</sup> | 0% | 0 | 128 | 7290 | 298 | 50 | [63] |
| +Höltje-CHOL-13<sup>64,65</sup> | 50% | 64 | 64 | 10314 | 298 | 50 | [66] |
| CHARMM36<sup>50</sup>/TIP3P<sup>51</sup> | 0% | 0 | 200 | 9000 | 310 | 500 | [67] |
| +CHARMM36<sup>68</sup> | 50% | 200 | 200 | 18000 | 310 | 500 | [69] |
| MacRog<sup>53</sup>/TIP3P<sup>54</sup> | 0% | 0 | 128 | 6400 | 310 | 500 | [70] |
| +MacRog<sup>53</sup> | 50% | 64 | 64 | 6400 | 310 | 500 | [70] |
| Slipids<sup>58</sup>/TIP3P<sup>54</sup> | 0% | 0 | 200 | 9000 | 310 | 500 | [59] |
| +Slipids<sup>71</sup> | 50% | 200 | 200 | 18000 | 310 | 500 | [59] |

<sup>a</sup>Bilayer cholesterol content (mol %).
<sup>b</sup>Number of cholesterol molecules.
<sup>c</sup>Number of POPC molecules.
<sup>d</sup>Number of water molecules.
<sup>e</sup>Simulation temperature.
<sup>f</sup>Trajectory length used for analysis.
<sup>g</sup>Reference for the openly available simulation files.

**Table 3:**
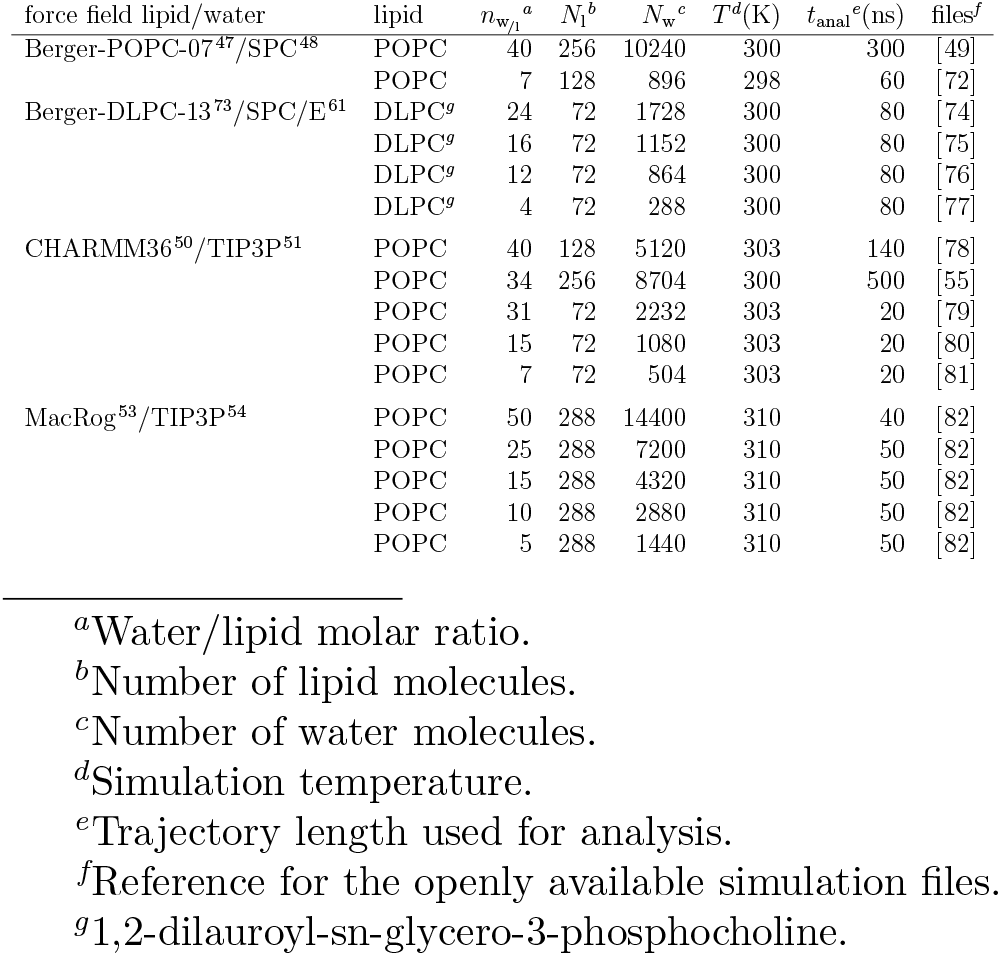
Analyzed open-access MD trajectories of PC lipid bilayers under varying hydration level.

The trajectories were analyzed using in-house scripts. These are available on GitHub (github.com/hsantila/Corrtimes/tree/master/teff_analysis), along with a Python notebook outlining an example analysis run. To enable automated analysis of several force fields with differing atom naming conventions, we used the mapping scheme developed within the NMRlipids Project to automatically recognise the atoms and bonds of interest for each trajectory.

After downloading the necessary files from Zenodo, we processed the trajectory with Gro-macs `gmx trjconv` to make the molecules whole; that is, we made sure that for each covalent bond the partaking atoms are from the same periodic image of the molecule. For the united atom Berger model, hydrogens were added using the Gromacs 4.0.2 tool `g_protonate`. We then calculated the *S*_CH_ (Eq. (3)) with the OrderParameter.py script that uses the MDanalysis^83,84^ Python library. The C–H bond correlation functions *g*(*τ*) (Eq. (1)) were calculated with Gromacs 5.1.4^85^ `gmx rotacf` (note that on MD timescales *g*_s_ = 1 so that *g* = *g*_f_) after which the *S*_CH_ were used to normalize the *g*_f_ to obtain the reduced and normalized correlation function

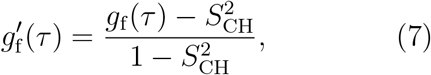

that is, the integrand in Eq. (4).

The effective correlation times *τ*_e_ were then calculated by integrating 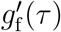 from *τ* = 0 until *τ* = *t*_0_. Here, *t*_0_ is the first time point at which 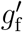 reached zero: 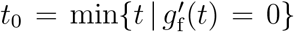. If 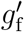 did not reach zero within *t*_anal_/2, the *τ*_e_ was not determined, and we report only its upper and lower estimates.

To quantify the error on *τ*_e_, we first estimate the error on 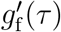, where we account for two sources of uncertainty: *g*_f_(*τ*) and 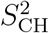. Performing linear error propagation on Eq. (7) gives

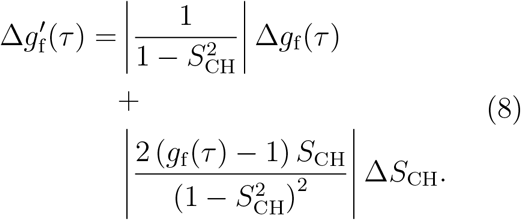

Here the Δ*S*_CH_ was determined as the standard error of the mean of the *S*_CH_ over the *N*_l_ individual lipids in the system. ^32^ Similarly, we quantified the error on *g*_f_ (*τ*) by first determining the correlation function 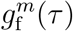 for each individual lipid *m* over the whole trajectory, and then obtaining the error estimate Δ*g*_f_ (*τ*) as the standard error of the mean over the *N*_l_ lipids. Importantly, this gives an uncertainty estimate for *g*_f_ (*τ*) at each time point *τ*.

To obtain the lower bound on *τ*_e_, we integrate the function 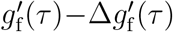 over time from *τ* = 0 until *τ* = *t*_l_. Here

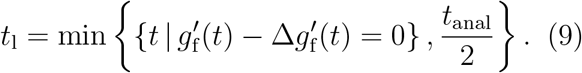

That is, *t*_l_ equals the first time point at which the lower error estimate of 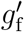 reached zero; or *t*_l_ = *t*_anal_/2, if zero was not reached before that point.

To obtain the upper error estimate on *τ*_e_, we first integrate the function 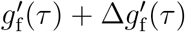 over time from *τ* = 0 until *t*_u_ = min {*t*_0_, *t*_anal_/2}. Note, however, that this is not yet sufficient, because there could be slow processes that the simulation was not able to see. Although these would contribute to *τ*_e_ with a low weight, their contribution over long times could still add up to a sizable effect on *τ*_e_. That said, it is feasible to assume (see Fig. 1A) that there are no longer-time contributions to *g*_f_ than something that decays with a time constant of 10^-6^ s. We use this as our worst case estimate to assess the upper bound for *τ*_e_, that is, we assume that all the decay of *g*_f_ from the time point *t*_u_ onwards comes solely from this hypothetical slowest process that decays with a time constant of 10^-6^ s. The additional contribution to the upper bound for *τ*_e_ then reads

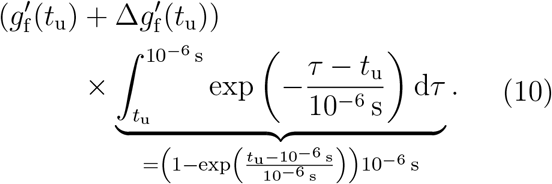

The *R*_1_ rates were calculated using Eq. (5). The spectral density *j* (*ω*) was obtained from the normalized correlation function 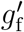 by fitting it with a sum of 61 exponentials

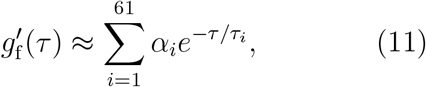

with logarithmically spaced time-scales *τ_i_* ranging from 1 ps to 1 μs, and then calculating the spectral density of this fit based on the Fourier transformation^41^

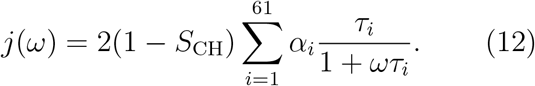

The *R*_1_ rate of a given C–H pair was first calculated separately for each lipid *m* (using Eq. (5) with *N*_H_ = 1, and *j*^m^(*ω*) obtained for the normalized correlation function 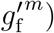. The resulting *N*_l_ measurements per C–H pair were then assumed independent: Their mean gave the *R*_1_ rate of the C–H pair, and standard error of the mean its uncertainty. The total *R*_1_ rate of a given carbon was obtained as a sum of the *R*_1_ rates of its C–H pairs. When several carbons contribute to a single experimental *R*_1_ rate due to the overlapping peaks (for example in C2 carbon in the acyl chains and the *γ* carbons), the *R*_1_ from simulations was obtained as an average over carbons with overlapping peaks. The segment-wise error estimates were obtained by standard error propagation, starting from the uncertainties of the *R*_1_ rates of the C–H pairs.

To gain some qualitative insight on the time scales at which the main contributions to the *R*_1_ rates arise, we also calculated ‘cumulative’ *R*_1_ rates, *R*_1_(*τ*), which contained those terms of the sum in Eq. (12) for which *τ_i_* < *τ*. Note that here the 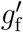 averaged over lipids was used; therefore, the ‘cumulative’ *R*_1_(*τ* → ∞) does not necessarily have exactly the same numerical value as the actual *R*_1_.

Finally, we note that the fit of Eq. (11) provides an alternative to estimating *τ*_e_, because

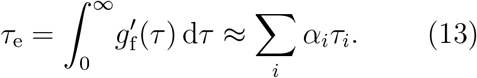

When the simulation trajectory is not long enough for the correlation function to reach the plateau, integrating 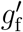 gives a lower bound estimate for *τ*_e_, while the sum of Eq. (13) includes also (some) contribution from the longer-time components via the fitting process. However, in practice the fit is often highly unreliable in depicting the long tails of the correlation function, and thus we chose to quantify *τ*_e_ using the area under 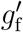, and estimate its uncertainty as detailed above.

## 3 Results and Discussion

Using open-access MD simulation trajectories, we benchmark phospholipid conformational dynamics in six MD force fields. We start with pure POPC bilayers in their liquid crystalline fully hydrated state (see Table 1 for simulation details and Fig. 3 for the data), and then proceed to check the changes in dynamics when cholesterol is added to the bilayer (Table 2 and Fig. 5) and when the hydration level is reduced (Table 3 and Fig. 6). Our yardsticks are the effective correlations times *τ*_e_ (Eq. (4)) and the *R*_1_ rates (Eq. (5)) measured at 125 MHz ^13^C (500 MHz ^1^H) Larmor frequency; an MD model with correct rotational dynamics in a window around ~1ns will match the experimental *R*_1_ rates, whereas the *τ*_e_ reflect all the sub-*μ*s time scales (Fig. 1).

**Figure 3:**
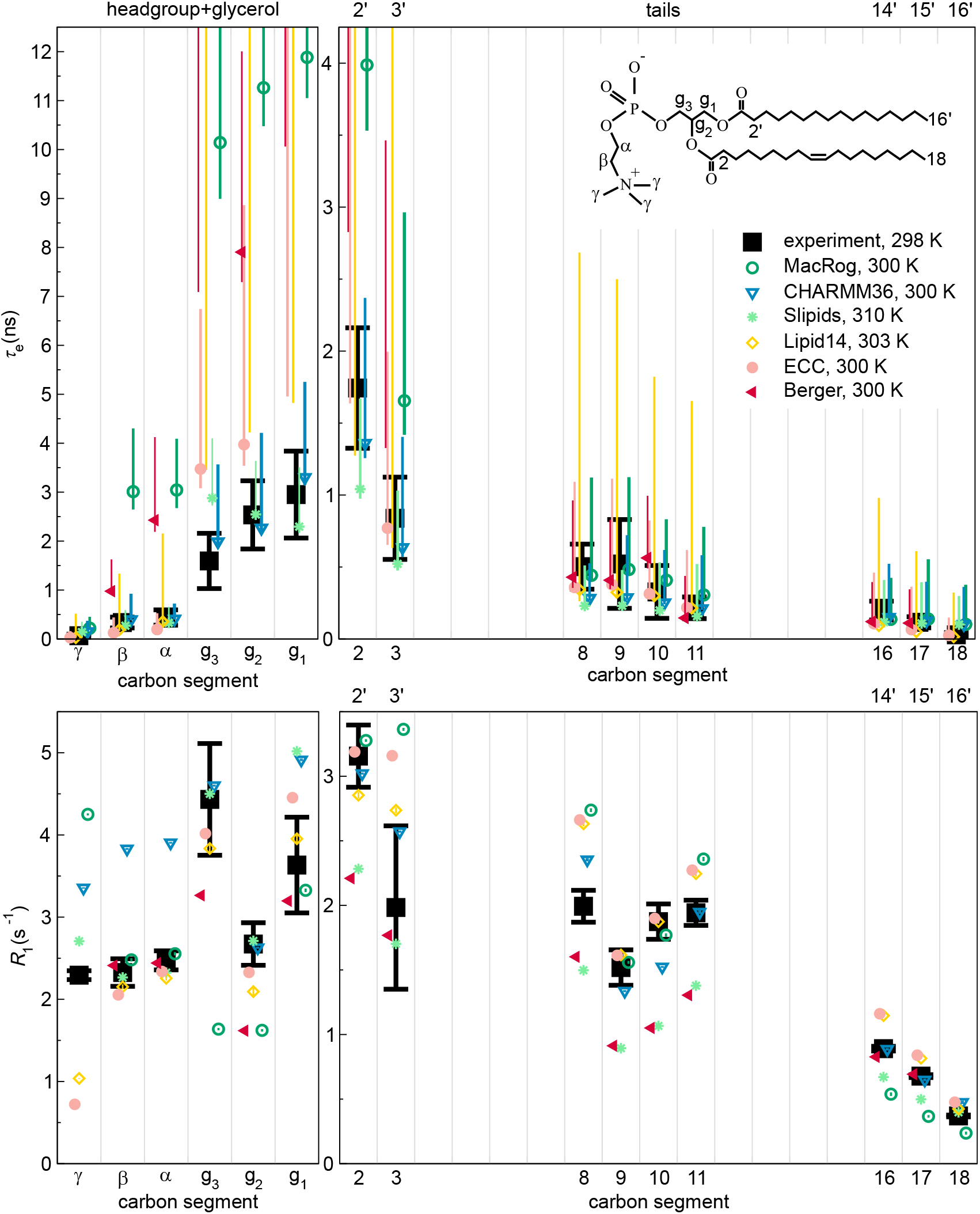
Effective correlation times (*τ*_e_, top) and *R*_1_ rates (bottom) in experiments^39^ (black) and MD simulations (colored) of POPC bilayers in *L_α_* phase under full hydration. Inset shows the POPC chemical structure and carbon segment labeling. Each plotted value contains contributions from all the hydrogens within its carbon segment; the data for segments 8–11 are only from the sn-2 (oleoyl) chain, whereas the (experimentally non-resolved) contributions of both tails are included for segments 2–3 (2’–3’ in the sn-1 chain) and 16–18 (14’–16’)· Simulation results are only shown for the segments for which experimental data was available. For *τ*_e_, a simulation data point indicates the average over C–H bonds; however, if *τ*_e_ could not be determined for all bonds, only the error bar (extending from the mean of the lower to the mean of the upper error estimates) is shown. The Berger data for segments *γ*, C18, and C16’ are left out, as the protonation algorithm used to construct the hydrogens post-simulation in united atom models does not preserve the methyl C–H bond dynamics. Table 1 provides further simulation details, while information on the experiments is available at Ref. 39.

### Pure POPC at full hydration: Slipids and CHARMM36 reproduce *τ*_e_ excellently

The top panels of Fig. 3 compare the effective correlation times *τ*_e_ obtained for fully hydrated POPC bilayers in experiments (black) and in six different MD force fields (color). We see that—as implied by the discussion leading to Eq. (10)—sub-*μ*s MD simulations typically lead to asymmetric error bars on *τ*_e_; if these openaccess trajectories were extended, the *τ*_e_ values would more likely increase than decrease. Qualitatively, every force field captures the general shape of the *τ*_e_ profile: Dynamics slows down towards the glycerol backbone in both the head-group and the tails.

Quantitatively, most MD simulations tend to produce too slow dynamics in the glycerol region (Fig. 3). This is consistent with previous results for the Berger model, ^41^ and with the insufficient conformational sampling of glycerol backbone torsions observed in 500-ns-long CHARMMc32b2^86,87^ simulations of a PC lipid. ^88^

The best overall *τ*_e_-performance is seen in Slipids and in particular CHARMM36 (Fig. 3). This is in line with CHARMM36 reproducing the most realistic conformational ensembles for the headgroup and glycerol backbone among the MD simulation force fields benchmarked here. ^32,34^ Indeed, it is important to keep in mind that the conformational ensembles greatly differ between force fields and are not exactly correct in any of them. ^32,34^ Consequently, the calculated *τ*_e_ times and *R*_1_ rates depict the dynamics of sampling a somewhat different and incorrect phase space for each model. To this end, we try to avoid overly detailed discussion on the models and rather concentrate on common and qualitative trends. That said, there are a few carbon segments in the data for which the experimental order parameters, *R*_1_, and *τ*_e_ are all (almost) reproduced by simulations, suggesting that both the conformational ensemble and the dynamics are correctly captured by MD in these cases. For example, Slipids performs well at the *β* and *α*, and CHARMM36 at the *g*_3_, *g*_2_, and C2 segments. These are, however, exceptions.

### An excellent *τ*_e_ may be accompanied by a poor *R*_1_, or *vice versa*

The lower panels of Fig. 3 compare the experimental and simulated *R*_1_ rates under the same conditions that were used for the *τ*_e_ above. Notably, there are several instances where the *R*_1_ comparison distinctly differs from what was seen for *τ*_e_.

There are cases where a matching *R*_1_ is accompanied by a larger-than-experimental *τ*_e_. MacRog for the *β, α*, and *g*_1_ segments provides a prominent example of this. Such a combination suggests that MD has the correct relative weight of 1-ns-scale dynamics, but has too slow long-time dynamics.

There are also cases where *τ*_e_ matches experiments, but *R*_1_ does not, such as the *β and α* segments in CHARMM36. Therein a cancellation of errors occurs in *τ*_e_: The overestimation of the relative weight of 1-ns-scale dynamics is compensated by wrong dynamics at the other time scales. As CHARMM36 overall performs rather well for C–H bond order parameters, *R*_1_, and *τ*_e_, we proceed to study this shortcoming on the headgroup *R*_1_ rates in some more detail.

### Conformational dynamics of PC headgroup segments in MD

Figure 4A zooms in on the headgroup (*γ, β, α*) segments, whose *τ*_e_ were not clearly visible on the scale of Fig. 3. We see that for *γ*, no force field provides both *τ*_e_ and *R*_1_, but Slipids comes closest. For *β* and *α*, Slipids captures both measurables near perfectly. In other words, among the benchmarked force fields Slipids gives the most realistic description of the conformational dynamics in the headgroup region. CHARMM36, e.g., overestimates (*R*_1_) the relative weight of timescales around ~1 ns.

**Figure 4:**
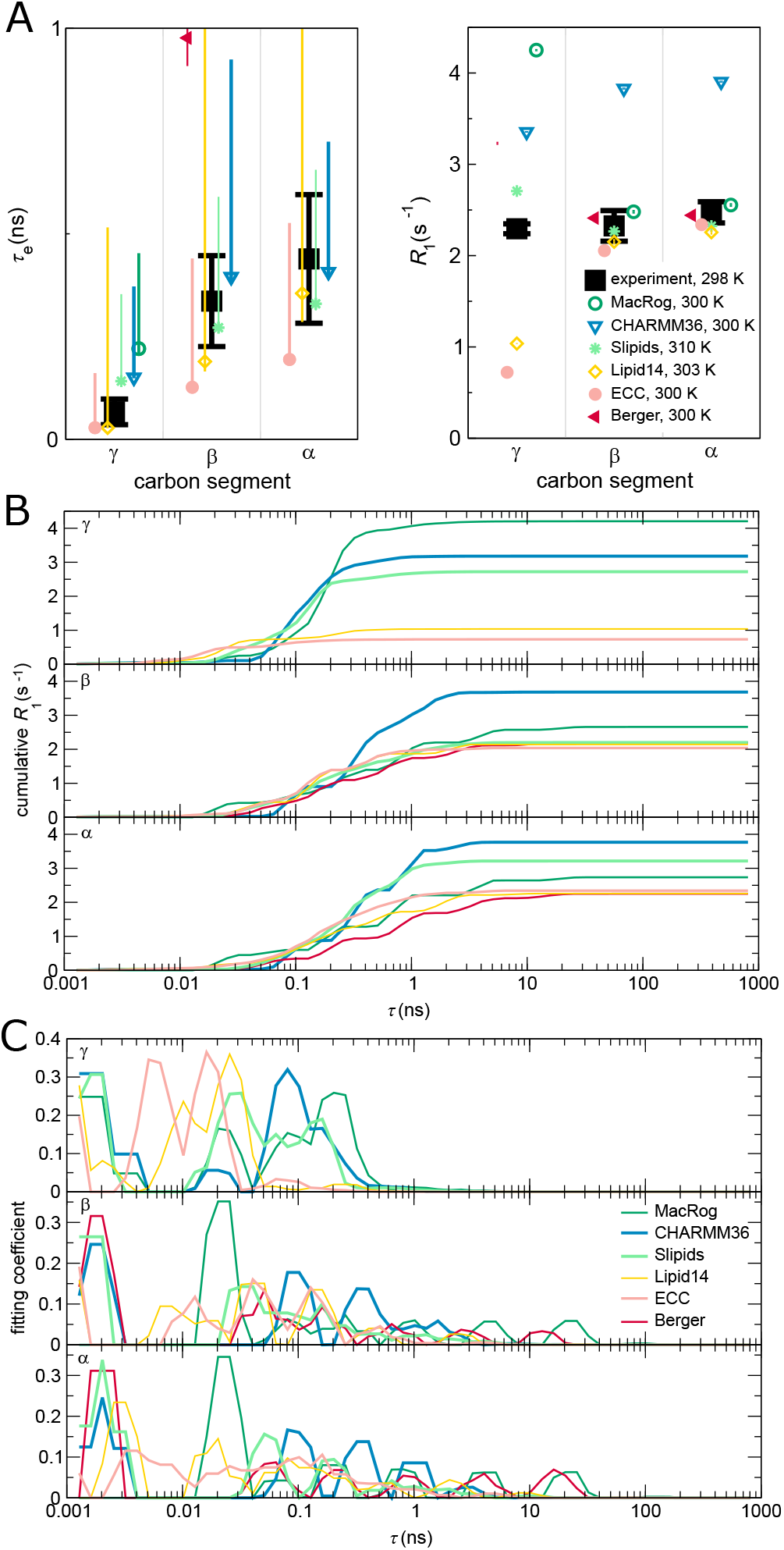
Contributions to the dynamics of the headgroup segments. (A) Zoom on the head-group *τ*_e_ (left panel) and *R*_1_ (right). (B) ‘Cumulative’ *R*_1_ (*τ*) of the *γ* (top panel), *β* (middle), and *α* (bottom) segments. *R*_1_(*τ*) is obtained, as detailed in Methods, by including in the sum of Eq. (12) only terms with *τ_i_* < *τ*. Consequently, at *τ* → ∞ the *R*_1_(τ) approaches the actual *R*_1_. (C) Prefactor weights *α_i_* from Eq. (11) of *γ* (top), *β* (middle), and *α* (bottom). Note that panels B and C show a sliding average over 3 neighboring data points.

To investigate closer how the differences between force fields arise, Fig. 4B shows the ‘cumulative’ *R*_1_(*τ*), where the ranges of steepest increase indicate time scales that most strongly contribute to *R*_1_ rates.

For the *γ* segment, Fig. 4B shows that for models that overestimate the *R*_1_ rate (MacRog, CHARMM36, and Slipids, see Fig. 4A) the major contribution to *R*_1_ arises at *τ* > 50 ps, whereas for models that underestimate *R*_1_ (Lipid14 and ECC) the major contribution comes from *τ* < 50 ps. This also manifests in the distribution of fitting weights (*α_i_* in Eq. (11)) in Fig. 4C: The later non-zero weights occur, the larger is the resulting *R*_1_ of *γ*.

For the *β* and *α* segments, Fig. 4B shows that the main contribution to *R*_1_ rates arises from processes between 100 ps and 1 ns. CHARMM36 has the largest relative weights of all models in this window (Fig. 4C), which explains its overestimation of *R*_1_ of *β* and *α*. All the other models have *R*_1_ rates close to experiments, but only Slipids simultaneously gives also the *τ*_e_ correctly. Notably, Slipids has its largest weights at *τ* < 100 ps. Indeed, the considerable weights at short (< 10 ps) time scales in all models except MacRog and at long (> 10 ns) time scales in MacRog and Berger hardly manifest in *R*_1_. However, the latter contribute heavily to *τ*_e_, which is thus considerably overestimated by MacRog and Berger (Fig. 3).

It would be highly interesting to identify the origins of the observed artificial timescales, particularly for the 0.1–1 ns window over-presented in CHARMM36, and propose how to correct those in the simulation models. After all, it is known that the *R*_1_ rates of mono- and disaccharides^89^ and proteins^90^ in solution agree satisfactorily with experiments when the artificially low viscosity of TIP3P water is accounted for by a simple scaling. Viscosity at the bilayer–water interface, however, remains an open question— although one which a careful comparison between spin relaxation rates of lipid headgroups in simulations and experiments might be able to answer. Nevertheless, we refrain from further analysis here, as the connection between the fitted correlation times and the correlation times of distinct motional processes, such as dihedral rotations and lipid wobbling, turns out to be highly non-trivial.

### Effect of cholesterol

An essential component in cell membranes, cholesterol has various biological functions. It is well known to order the acyl chains in lipid bilayers, but its effect on the headgroup is more controversial. ^65,91^ For example, it has been proposed that lipid headgroups reorganize to shield cholesterol from water. ^91^ However, while acyl chains do substantially order, NMR experiments show no significant conformational changes in the headgroup upon addition of even 50% of cholesterol—which suggests that the tail and head regions behave essentially independently. ^32,65^ In principle, the head-groups could shield cholesterol from water even without changing their conformational ensemble: By reorienting only laterally on top of the cholesterol. In this case, one would expect the rotational dynamics of headgroup segments to change when cholesterol is added.

Top panels of Fig. 5A depict the experimental effective correlation times *τ*_e_ in pure POPC bilayers and in bilayers containing 50% cholesterol. The *τ*_e_ at the glycerol backbone slow down markedly when cholesterol is added. Tail segment dynamics slows down too, most notably close to the glycerol backbone. In stark contrast, *τ*_e_ of the headgroup segments (*γ, β, α*) remain unaffected. Furthermore, cholesterol induces no measurable change in the headgroup *β* and *α* segment dynamics at short (~1ns) time scales, as demonstrated by the experimental *R*_1_ rates (Fig. 5A, bottom panels). That said, there is a small but measurable impact on *R*_1_ at *γ*. In summary, these experimental findings support the idea^39^ that the acyl chains and the headgroup can respond—in agreement with the relative uncoupling of the PC head and tails reported in simulations^92^—almost independently to changes in conditions and composition.

**Figure 5:**
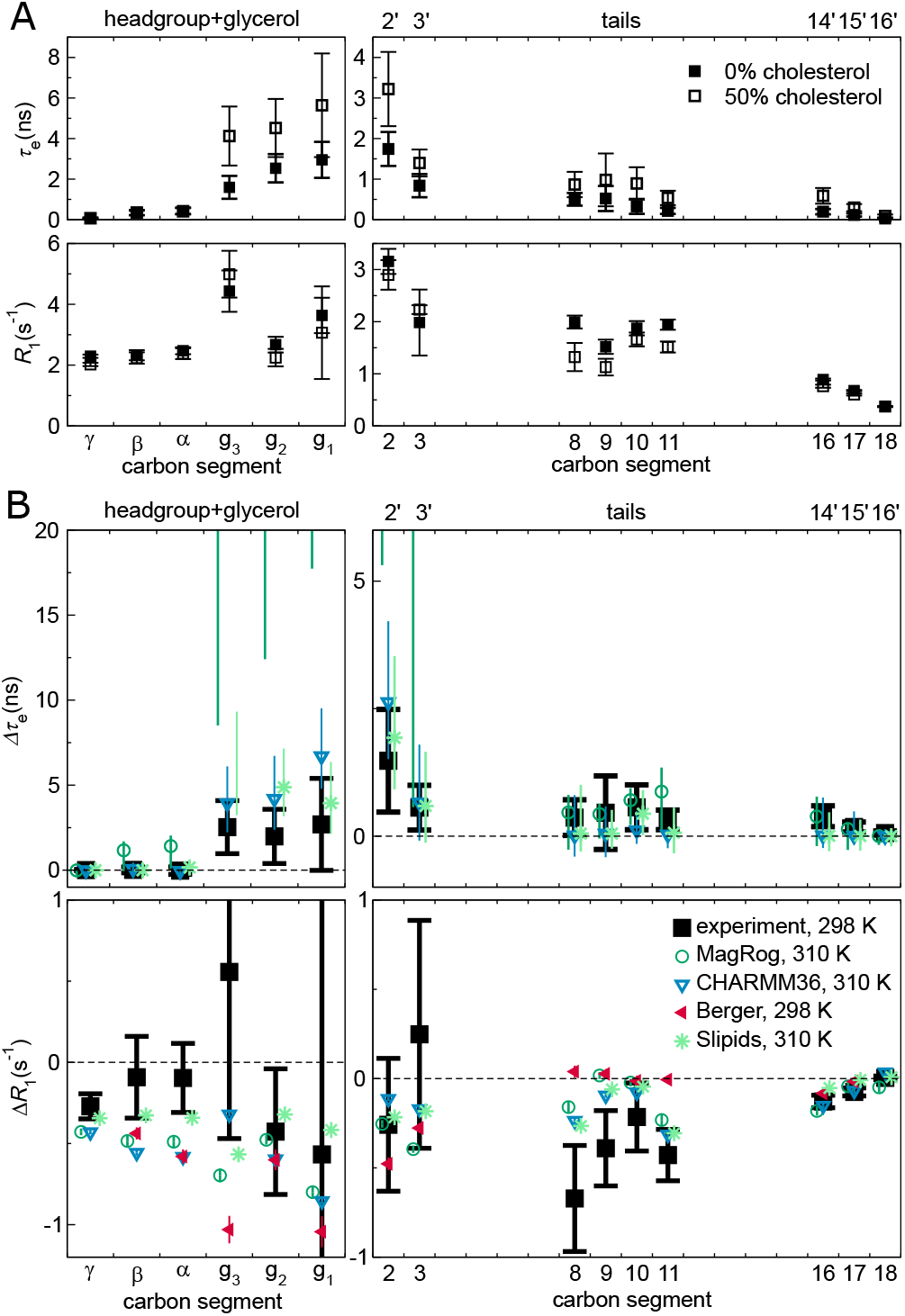
Effect of cholesterol on POPC conformational dynamics. (A) Experimental effective correlation times *τ*_e_ (top panels) and *R*_1_ rates (bottom) in 100/0 and 50/50 POPC/cholesterol bilayers at full hydration, see Ref. 39 for further details. (B) The change in *τ*_e_ (Δ*τ*_e_, top panels) and *R*_1_ (Δ*R*_1_, bottom), in NMR (black) and MD (color), when bilayer composition changes from pure POPC to 50% cholesterol. Error estimates for the simulated Δ*τ*_e_ are the maximal possible based on the errors at 0% and 50% cholesterol; for other data regular error propagation is used. The Berger Δ*τ*_e_ is not shown, because the available open-access trajectories were too short to determine meaningful error estimates. Table 2 provides further simulation details; for segment labeling, see Fig. 3.

All four benchmarked force fields (Fig. 5B) qualitatively reproduce the experimental increase in *τ*_e_: Slipids and CHARMM36 give rather decent magnitude estimates, while MacRog grossly overestimates the slowdown of glycerol, C2, and C3 segments. Notably, MacRog appears to predict slowdown also for the headgroup (*β* and *α*), for which experiments detect no change. Note that while CHARMM36 correctly shows no change in *τ*_e_ of the *γ, β*, and *α* segments, it does predict an erroneous Δ*R*_1_ for all three, indicating some inaccuracies in the headgroup rotational dynamics. Such inaccuracies might be reflected in the recent findings ^93^ (obtained using CHARMM36) that the head-groups of PCs neighboring (within 6.6 Å) a lone cholesterol spend more time on top of the said cholesterol than elsewhere. Interestingly, the tail Δ*R*_1_ seem to be qualitatively reproduced by all three all-atom force fields, whereas Berger fails to capture the trend at the oleoyl double bond. All these findings are in line with the general picture obtained from C–H bond order parameters: ^38^ MD simulations capture the changes in acyl chain region rather well, but changes in and near the glycerol backbone region can be overestimated. Of the benchmarked force fields, CHARMM36 appears most realistic in reproducing the effects of cholesterol on the glycerol backbone—and Slipids on the PC headgroup—conformational dynamics.

### Effect of drying

Understanding the impact of dehydration on the structure and dynamics of lipid bilayers is of considerable biological interest. Dehydrated states are found, e.g., in skin tissue. Most prominently, the process of membrane fusion is always preceeded by removal of water between the approaching surfaces, and thus the dehydration-imposed changes can considerably affect fusion characteristics, such as its rate.

Figure 6A shows how a mild dehydration affects C–H bond dynamics in the PC head-group and glycerol backbone; the plot compares the experimental effective correlation times *τ*_e_ measured for POPC at full hydration and for DMPC (1,2-dimyristoyl-sn-glycero-3-phosphocholine) at 13 waters per lipid. The *τ*_e_ are the same within experimental accuracy, which suggests two conclusions. Firstly, the headgroup (*γ, β, α*) *τ*_e_ are rather insensitive to the chemical identities of the tails. This is analogous to what was seen experimentally when adding cholesterol (Fig. 5A): Structural changes in the tail and glycerol regions do not (need to) affect the headgroup dynamics. Secondly, a mild dehydration does not alter the *τ*_e_ in the headgroup and glycerol regions.

**Figure 6:**
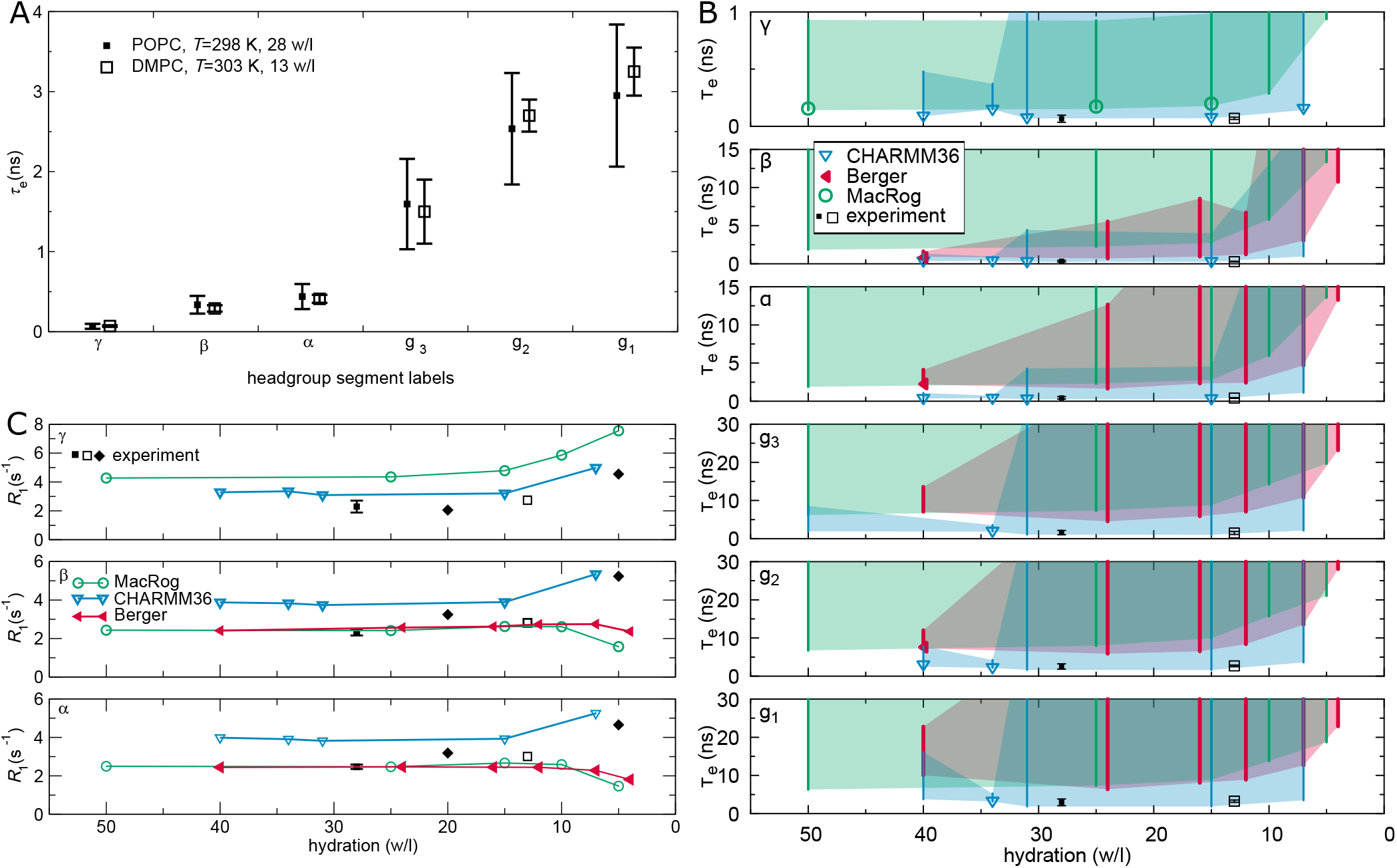
Effect of drying on PC headgroup and glycerol backbone conformational dynamics. (A) Experimental effective correlation times *τ*_e_ for DMPC at low hydration (from Ref. 42) do not significantly differ from the *τ*_e_ for POPC at full hydration (from Ref. 39). (B) Calculated *τ*_e_ for POPC at decreasing hydration in three MD models. Symbols indicate the mean of segment hydrogens if *τ*_e_ could be determined for all of them; otherwise, only the error bar (extending from the mean of the lower to the mean of the upper uncertainty estimates) is drawn. The area limited by the error bars shaded for visualization. Note that four Berger data points (24, 16, 12, and 4 w/l) are from DLPC. (C) ^13^C-NMR *R*_1_ rates (at *ω*_C_ = 125 MHz) of the PC headgroup segments in experiments and simulations: Experiments indicate an increasing trend upon dehydration. Experimental POPC (*T* = 298K) data at 28 w/l is from Ref. 39 (■), POPC (298K) at 20 and 5w/l from Ref. 43 (♦), and DMPC (303K) at 13 w/l from Ref. 42 (□). See Table 3 for simulation details.

Figure 6B shows the effects of dehydration in three MD models. Combination of the unrealistically slow dynamics, especially in the glycerol backbone (Fig. 3), and the relatively short lengths of the available open-access trajectories (Table 3) led to large uncertainty estimates; thus we only point out qualitative trends here. For all headgroup and glycerol segments, the simulated *τ*_e_ indicate slowdown upon dehydration. This is manifested in the increase in the magnitude of the error estimate (cf. the Berger data for *β* and *α*) as well as in the increase of the lower limit of the error. For CHARMM36 the lower error estimates stay almost constant all the way until 7w/l, whereas for Berger and MacRog they hint that a retardation of dynamics starts already between 15 and 10 w/l.

These simulational findings suggest that experiments reducing hydration levels below 10w/l would also show an increase in *τ*_e_. This prediction is in line with the exponential slowdown of the headgroup conformational dynamics upon dehydration that was indicated by ^2^H-NMR *R*_1_ measurements of DOPC bilayers: *R*_1_ ~ exp(−*n*_w/l_/4).^94^ The slowdown was attributed to the reduced effective volume available for the headgroup ^94^ as it tilts towards the membrane upon dehydration; such tilt is observed via changes of the lipid headgroup order parameters, ^95^ and is qualitatively reproduced by all the simulation models. ^32^

Figure 6C shows a collection of experimental ^13^C-NMR *R*_1_ rates for the headgroup segments at different water contents; in addition to the full hydration POPC data from Fig. 3, DMPC at 13w/l,^42^ and POPC at 20 and 5w/l^43^ are shown. Experimentally, an increasing trend with decreasing hydration is observed for all three segments, indicating changes of head-group dynamics at short (~1ns) time scales. Interestingly, only CHARMM36 captures this, whereas Berger and MacRog give decreasing *R*_1_ rates for *β* and *α*.

The slowdown characteristics discussed here are of significance not only for computational studies of intermembrane interactions, such as fusion, but also when simulating a bilayer (stack) under low hydration: Slower dynamics require longer simulation times for equilibration, for reliably quantifying the properties of the bilayers, and for observing rare events.

## 4 Conclusions

We have here demonstrated that open access databanks of MD trajectories enable the creation of new scientific information without running a single new simulation. More specifically, we have benchmarked (against published NMR data^39,41-43^) the conformational dynamics of a wide range of phosphatidylcholine MD models using existing open-access trajectories from the Zenodo repository, in particular those belonging to the NMRlipids Databank (zenodo.org/communities/nmrlipids).

We found that every MD model captures the ^13^C-NMR effective correlation time (*τ*_e_) profile of POPC qualitatively, but that most are prone to too slow dynamics of the glycerol backbone C–H bonds (Fig. 3). While no force field perfectly reproduces all the experimental data, CHARMM36 and Slipids have overall impressive *τ*_e_. This is a particularly exciting finding concerning CHARMM36, as it is also known to reproduce quite well the experimental conformational ensemble. ^32^ That said, we do find that CHARMM36 struggles with the balance of dynamics in the headgroup region: The *R*_1_ rates, sensitive for ~1-ns processes, are too high for the *γ, β*, and *α* segments (Fig. 4). In fact Slipids, which also reproduces the experimental headgroup order parameters, ^32^ appears to out-perform CHARMM36 when it comes to head-group conformational dynamics (Fig. 4).

Further, we found that when cholesterol is mixed into a POPC bilayer, MD qualitatively captures the slowdown of conformational dynamics in the tail and glycerol regions (Fig. 5). However, the benchmarked force fields overestimate the changes in the ~1-ns dynamics of the headgroup—except Slipids, which captures well the effects of cholesterol on PC headgroup conformational dynamics.

Finally, we found that upon reducing the water content below 10 waters per lipid, MD exhibits slowdown of headgroup and backbone dynamics in qualitative agreement with experimental data. That said, only CHARMM36 (but not Berger or MacRog) qualitatively captures the experimentally detected increase of *R*_1_ rates upon dehydration (Fig. 6).

While work is still needed in capturing even the correct phospholipid conformations, ^32^ realistic dynamics will be an essential part of developing MD into a true computational microscope. Here we gathered a set of published experimental ^13^C-NMR data on phosphatidylcholine conformational dynamics, and charted the typical features of the existing MD models against it, thus laying the foundation for further improvement of MD force fields. Importantly, our work demonstrates the potential of open-access MD trajectories in achieving such benchmarks at a reduced computational and labor cost—but it also highlights the challenges inherent in using such data: Not all system permutations might readily exist (here the dehydration data for Lipid14 and Slipids were lacking, see Fig. 6); the available sampling (simulation length and size) might vary, requiring extreme care with error estimation; and one has to remain aware of the subtle differences in the many simulation parameters (barostats, temperatures, characteristics of the MD engines,…). That said, it has not escaped our notice that a pool of well indexed and documented open-access data provides an ideal platform for automation, which in turn will facilitate faster progress in pinpointing the typical failures of the existing force fields, in identifying key differences in models describing chemical variations of the same molecule type (such as different lipid headgroups), and in developing better models through data-driven approaches.

## Acknowledgement

H. S. A. gratefully acknowledges the support from Osk. Huttunen Foundation, Finnish Academy of Science and Letters (Foundations’ Post Doc Pool), Instrumentarium Science Foundation, and the Alexander von Humboldt Foundation. O. H. S. O. acknowledges the Academy of Finland (315596, 319902) for financial support.

## Graphical TOC Entry

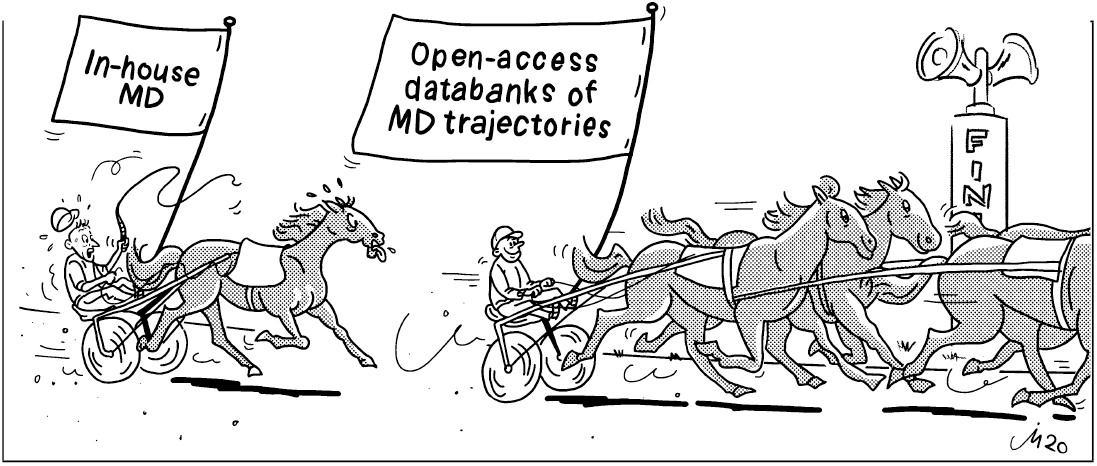

